# SARS-CoV-2 Neutralizing Antibodies After Bivalent vs. Monovalent Booster

**DOI:** 10.1101/2023.02.13.528341

**Authors:** Qian Wang, Anthony Bowen, Anthony R. Tam, Riccardo Valdez, Emily Stoneman, Ian A. Mellis, Aubree Gordon, Lihong Liu, David D. Ho

**Author notes:** Correspondence (A.G.), (L.L.), (D.D.H.). These authors contributed equally (Qian Wang, Anthony Bowen).

## Abstract

Bivalent mRNA vaccine boosters expressing Omicron BA.5 spike and ancestral D614G spike were introduced to attempt to boost waning antibody titers and broaden coverage against emerging SARS-CoV-2 lineages. Previous reports showed that peak serum neutralizing antibody (NAb) titers against SARS-CoV-2 variants following bivalent booster were similar to peak titers following monovalent booster. It remains unknown whether these antibody responses would diverge over time. We assessed serum virus-neutralizing titers in 41 participants who received three monovalent mRNA vaccine doses followed by bivalent booster, monovalent booster, or BA.5 breakthrough infection at one month and three months after the last vaccine dose or breakthrough infection using pseudovirus neutralization assays against D614G and Omicron subvariants (BA.2, BA.5, BQ.1.1, and XBB.1.5). There was no significant difference at one month and three months post-booster for the two booster cohorts. BA.5 breakthrough patients exhibited significantly higher NAb titers at three months against all Omicron subvariants tested compared against monovalent and bivalent booster cohorts. There was a 2-fold drop in mean NAb titers in the booster cohorts between one and three month time points, but no discernible waning of titers in the BA.5 breakthrough cohort over the same period. Our results suggest that NAb titers after boosting with one dose of bivalent mRNA vaccine are not higher than boosting with monovalent vaccine. Perhaps inclusion of D614G spike in the bivalent booster exacerbates the challenge posed by immunological imprinting. Hope remains that a second bivalent booster could induce superior NAb responses against emerging variants.

## Main text

The severe acute respiratory coronavirus 2 (SARS-CoV-2) Omicron lineage continues to proliferate and evolve, leading to subvariants adept at evading antibody responses^1^. Bivalent mRNA vaccines expressing both the Omicron BA.5 spike and the ancestral D614G spike were introduced in August 2022 with the goal of boosting waning antibody titers and broadening coverage against emerging SARS-CoV-2 lineages^2^. However, we and others recently reported that peak serum neutralizing antibody (NAb) titers against SARS-CoV-2 variants following a bivalent vaccine booster were similar to peak titers following a monovalent booster^3,4^. It remains unknown whether these antibody responses would diverge over time.

We addressed this question by assessing serum virus-neutralizing titers in 41 participants who received three monovalent mRNA vaccines followed by a bivalent booster, a monovalent booster, or a BA.5 breakthrough infection (**Table S1**). We collected serum samples at nearly one month and approximately three months following the last vaccine dose or breakthrough infection (**Table S2)** and then determined their NAb titers using a pseudovirus neutralization assay^4^ against the ancestral D614G strain and a panel of Omicron subvariants: BA.2, BA.5, BQ.1.1, and XBB.1.5 (see Supplementary Appendix). Participants who received a monovalent booster were older (mean 55.3 years) than those who received a bivalent booster (mean 37.8 years) or those who had a breakthrough infection (mean 44.0 years) (**Table S2**).

Each cohort exhibited the highest serum NAb titers (ID_50_) against D614G and substantially lower titers against the latest Omicron subvariants. Consistent with our prior study^4^, there was no significant difference at nearly one month post last booster for the two vaccine cohorts (**Figure 1A**). At approximately three months post, there were again no statistical difference between the two groups (**Figure 1B**), although there was a trend favoring the bivalent group (1.4 to 1.6-fold). The BA.5 breakthrough cohort exhibited significantly higher NAb titers at three months against all Omicron subvariants tested when compared to both monovalent and bivalent booster cohorts (**Figure 1B**). An approximately 2-fold drop in mean NAb titers was noted for the vaccine cohorts over the follow-up period of ∼2 months (**Figure 1C**), consistent with recent findings^5^. Interestingly, there was no discernible waning of antibody responses in the BA.5 breakthrough infection cases over the same observation period.

**Figure 1.**
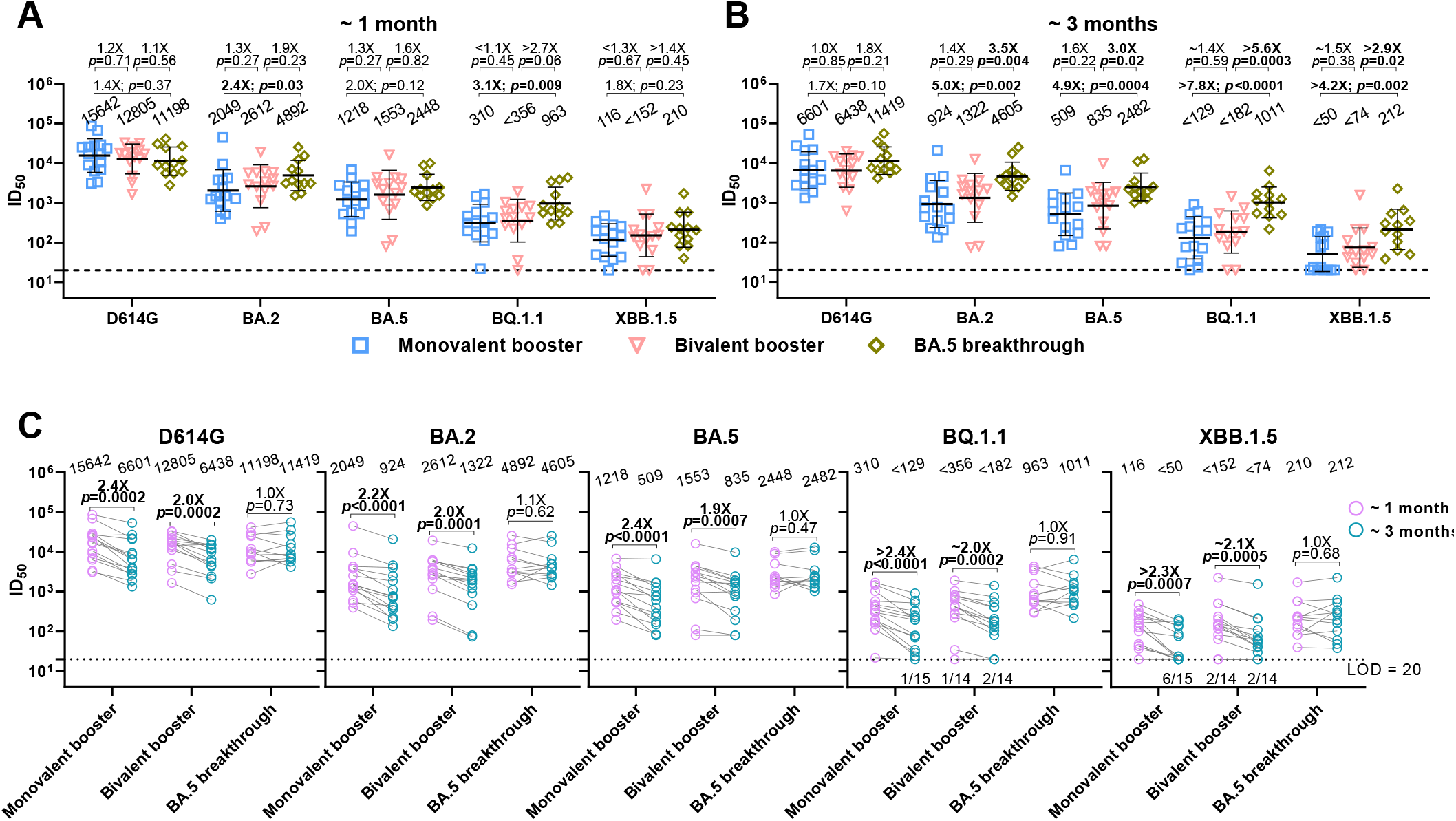
SARS-CoV-2 NAb responses following bivalent or monovalent booster vaccination or BA.5 breakthrough infection. **(A)** Peak serum neutralizing ID_50_ titers at nearly one month from “Monovalent booster”, “Bivalent booster”, and “BA.5 breakthrough” cohorts. **(B)** Serum neutralizing ID_50_ titers at ∼3 months for the three cohorts. **(C)** Paired serum neutralizing ID_50_ titers are shown at ∼1 and ∼3 months following most recent vaccination or breakthrough infection. Values above symbols denote the geometric mean ID_50_ titer (GMT) for each cohort. Comparisons show the fold change in GMT and were made by Mann-Whitney tests with *p*-values marked in bold if *p* < 0.05. The assay limit of detection (LOD) is 20 (dotted line) and the number of samples at or below the LOD is denoted above the x-axis.

These new results further strengthen our initial suggestion that boosting with the bivalent mRNA vaccines is not evidently better than boosting with the original monovalent vaccine as judged by serum SARS-CoV-2-neutralizing potency and breadth. Perhaps the inclusion of the ancestral spike in the bivalent vaccine exacerbates the challenge posed by immunological imprinting. Hope remains that a second bivalent vaccine booster could induce a superior NAb response against current and future viral variants.

## References

1. Wang Q, Iketani S, Li Z, et al. Alarming antibody evasion properties of rising SARS-CoV-2 BQ and XBB subvariants. Cell 2023;186:279–86 e8.

2. Coronavirus (COVID-19) Update: FDA Authorizes Moderna, Pfizer-BioNTech Bivalent COVID-19 Vaccines for Use as a Booster Dose. U.S. Food & Drug Administration, 2023. at https://www.fda.gov/news-events/press-announcements/coronavirus-covid-19-update-fda-authorizes-moderna-pfizer-biontech-bivalent-covid-19-vaccines-use.)

3. Collier AY, Miller J, Hachmann NP, et al. Immunogenicity of BA.5 Bivalent mRNA Vaccine Boosters. N Engl J Med 2023;388:565–7.

4. Wang Q, Bowen A, Valdez R, et al. Antibody Response to Omicron BA.4-BA.5 Bivalent Booster. N Engl J Med 2023;388:567–9.

5. Lasrado N, Collier AY, Miller J, et al. Waning Immunity Against XBB.1.5 Following Bivalent mRNA Boosters. bioRxiv 2023.

