## Supplemental information for "SARS-CoV-2 Neutralizing Antibodies After Bivalent vs. Monovalent Booster"

**Supplementary Methods**

*Clinical cohorts*

The sera analyzed in this study were obtained from three cohorts: "monovalent booster", “bivalent booster”, and “BA.5 breakthrough”. The first cohort comprised of individuals who received four doses of the monovalent COVID-19 mRNA vaccine, whereas the second cohort comprised of individuals who received three doses of the monovalent COVID-19 mRNA vaccine followed by one dose of the Pfizer or Moderna bivalent mRNA vaccine. The last cohort comprised of patients who had Omicron BA.5 breakthrough infection after three doses of the monovalent mRNA vaccine. Serum samples were evaluated using anti-nucleocapsid protein (NP) ELISA to determine prior SARS-CoV-2 infection status.

A subset of the sera was collected from the University of Michigan through the Immunity-Associated with SARS-CoV-2 Study (IASO), an ongoing cohort study that started in 2020[1]. All participants in the IASO study provided written informed consent, and the serum samples were collected under a protocol approved by the Institutional Review Board of the University of Michigan Medical School. Another subset of the vaccinee and breakthrough sera was collected at Columbia University Irving Medical Center, where all subjects provided written informed consent, and the serum collections were performed under protocols reviewed and approved by the Institutional Review Board of Columbia University. The detailed information for each case is provided in **Table S1**, with clinical information for the different study cohorts summarized in **Table S2**.

*Cell lines*

Vero-E6 cells (CRL-1586) and HEK293T cells (CRL-3216) were procured from the American Type Culture Collection (ATCC). The cells were cultured in Dulbecco's Modified Eagle Medium (DMEM) supplemented with 10% fetal bovine serum and 1% penicillin-streptomycin, and maintained in a 5% $CO_2$ atmosphere at 37°C.

*SARS-CoV-2 spike plasmids*

Plasmids encoding the spike protein of various SARS-CoV-2 variants, including D614G, BA.2, BA.4/5, and BQ.1.1 were previously generated[2-5]. The XBB.1.5 spike was constructed using the QuikChange II XL site-directed mutagenesis kit as per the manufacturer's instructions (Agilent). The sequences of all the constructs were verified by Sanger sequencing before use in experiments.

*Pseudovirus production*

SARS-CoV-2 variants were generated by replacing the native glycoprotein of vesicular stomatitis virus (VSV) with the SARS-CoV-2 variant spike protein[6]. Briefly, transfection of HEK293T cells with plasmid encoding the relevant spike protein was carried out using polyethyleneimine (PEI) at a concentration of 1 mg/mL. The transfected HEK 293T cells were then incubated at 37°C in a 5% $CO_2$ atmosphere for 24 hours, followed by infection with the VSV-G pseudotyped ΔG-luciferase (G*ΔG-luciferase, Kerafast). After a 2-hour incubation period at 37°C, the infected HEK293T cells were washed three times and cultured in fresh medium for an additional 24 hours under the same conditions. The collected supernatants were then centrifuged to remove any precipitates, aliquoted and stored at -80°C. To eliminate the presence of contaminating VSV-G pseudotyped ΔG-luciferase, the viral stock was pre-incubated with 20% I1 hybridoma (anti-VSV-G) supernatant (ATCC; CRL-2700) for 1 hour at 37°C prior to infecting target cells.

*Pseudovirus neutralization*

Before conducting the neutralization assay, the titers of all pseudoviruses were standardized to ensure consistent viral input. The heat-inactivated sera were then assayed in triplicate using 96-well plates and serially diluted four-fold in media starting from a 1:20 dilution. The pseudoviruses were added to the diluted sera and incubated for 1 hour at 37 °C. Control wells containing only virus were included on all plates. Vero-E6 cells were added to each well at a density of $4x10^4$ cells/well and incubated for 10 hours at 37°C with 5% $CO_2$. The cells were lysed, and luciferase activity was quantified using the Luciferase Assay System (Promega) and SoftMax Pro v.7.0.2 (Molecular Devices), following the manufacturer's instructions.

*Quantification and statistical analysis*

The 50% inhibitory dilution ($ID_{50}$) was determined using a five-parameter dose-response curve analysis in GraphPad Prism v.9.2. Unpaired groups were compared using two-tailed Mann-Whitney test and paired groups were compared using Wilcoxon matched-pairs signed rank test, both performed in GraphPad Prism v.9.2.

**Acknowledgements**

This study was supported financially by the NIH SARS-CoV-2 Assessment of Viral Evolution (SAVE) Program, as well as the NIAID, NIH (Contract Number 75N93019C00051) awarded to A.G. A.B. was supported in part by T32AI100852, Columbia Integrated Training Program in Infectious Disease Research. The authors express their gratitude to David Manthei, Carmen Gherasim, Victoria Blanc, Pamela Bennett-Baker, Savanna Sneeringer, Lauren Warsinske, Theresa Kowalski-Dobson, Alyssa Meyers, Zijin Chu, Hailey Kuiken, Lonnie Barnes, Ashley Eckard, Kathleen Lindsey, Dawson Davis, Aaron Rico, Casey Juntila, Daniel Raymond, Mayurika

Patel, and Nivea Vydiswaran from the IASO study team for their contribution in providing the serum samples.

### Author Contributions

L.L. and D.D.H. conceived the study. Q.W., A.R.T., I.A.M., and L.L. performed experiments. Q.W. managed the project. A.B., R.V., E.S. and A.G. collected serum samples. Q.W., A.B., L.L., and D.D.H. analyzed the results and wrote the manuscript. L.L. and D.D.H. directed and supervised the project. All authors reviewed and approved of the manuscript.

### Declaration of Interests

The authors declare potential conflicts of interest as follows: D.D.H. is a co-founder of TaiMed Biologics and RenBio, as well as a board director for Vicarious Surgical; he also serves as a consultant to WuXi Biologics, Brii Biosciences, and Veru; and he receives funding from Regeneron. Aubree Gordon is a member of a scientific advisory board for Janssen Pharmaceuticals. The remaining authors declare no competing interests.

### Table S1. Demographics of clinical cohorts

| Sample ID | Vaccine type and infection strain | Days V2-V3 | Days V3-V4 #(V3-I) | Days post last booster or infection* | | Confirmed COVID-19 | Age | Gender |
|---|---|---|---|---|---|---|---|---|
| | | | | Blood draw 1 | Blood draw 2 | | | |
| *Monovalent booster* | | | | | | | | |
| 1 | BNT162b2/BNT162b2/BNT162b2/BNT162b2 | 260 | 181 | 24 | 113 | No | 52 | Female |
| 2 | BNT162b2/BNT162b2/BNT162b2/BNT162b2 | 239 | 224 | 20 | 93 | No | 57 | Female |
| 3 | BNT162b2/BNT162b2/BNT162b2/BNT162b2 | 271 | 177 | 20 | 78 | No | 61 | Female |
| 4 | BNT162b2/BNT162b2/BNT162b2/BNT162b2 | 337 | 122 | 23 | 78 | No | 50 | Female |
| 5 | BNT162b2/BNT162b2/BNT162b2/BNT162b2 | 260 | 193 | 22 | 105 | No | 50 | Female |
| 6 | BNT162b2/BNT162b2/BNT162b2/BNT162b2 | 290 | 182 | 20 | 74 | No | 58 | Female |
| 7 | BNT162b2/BNT162b2/BNT162b2/BNT162b2 | 295 | 162 | 26 | 104 | No | 56 | Female |
| 8 | BNT162b2/BNT162b2/BNT162b2/BNT162b2 | 228 | 132 | 29 | 105 | No | 63 | Female |
| 9 | BNT162b2/BNT162b2/BNT162b2/BNT162b2 | 288 | 178 | 25 | 112 | No | 58 | Female |
| 10 | BNT162b2/BNT162b2/BNT162b2/BNT162b2 | 201 | 199 | 26 | 74 | No | 54 | Female |
| 11 | BNT162b2/BNT162b2/BNT162b2/BNT162b2 | 244 | 228 | 23 | 78 | No | 53 | Male |
| 12 | BNT162b2/BNT162b2/BNT162b2/BNT162b2 | 323 | 156 | 21 | 85 | No | 55 | Female |
| 13 | BNT162b2/BNT162b2/BNT162b2/BNT162b2 | 313 | 151 | 23 | 115 | No | 59 | Female |
| 14 | BNT162b2/BNT162b2/BNT162b2/BNT162b2 | 252 | 241 | 21 | 89 | No | 49 | Female |
| 15 | BNT162b2/BNT162b2/BNT162b2/BNT162b2 | 284 | 294 | 27 | 87 | No | 55 | Female |
| *Bivalent booster* | | | | | | | | |
| 16 | BNT162b2/BNT162b2/BNT162b2/Moderna Bivalent | 259 | 327 | 24 | 90 | No | 38 | Female |
| 17 | BNT162b2/BNT162b2/BNT162b2/Moderna Bivalent | 218 | 281 | 27 | 97 | No | 42 | Female |
| 18 | mRNA-1273/mRNA-1273/mRNA-1273/Moderna Bivalent | 205 | 294 | 24 | 95 | No | 36 | Male |
| 19 | BNT162b2/BNT162b2/BNT162b2/Pfizer Bivalent | 275 | 330 | 25 | 111 | No | 49 | Female |
| 20 | BNT162b2/BNT162b2/BNT162b2/Moderna Bivalent | 213 | 280 | 25 | 82 | No | 37 | Female |
| 21 | BNT162b2/BNT162b2/BNT162b2/Pfizer Bivalent | 256 | 346 | 26 | 98 | No | 45 | Male |
| 22 | BNT162b2/BNT162b2/mRNA-1273/Moderna Bivalent | 223 | 288 | 26 | 91 | No | 43 | Female |
| 23 | mRNA-1273/mRNA-1273/mRNA-1273/Moderna Bivalent | 211 | 279 | 29 | 97 | No | 32 | Female |
| 24 | BNT162b2/BNT162b2/BNT162b2/Pfizer Bivalent | 226 | 290 | 23 | 100 | No | 43 | Female |
| 25 | BNT162b2/BNT162b2/mRNA-1273/Moderna Bivalent | 266 | 321 | 27 | 90 | No | 36 | Female |
| 26 | BNT162b2/BNT162b2/BNT162b2/Moderna Bivalent | 254 | 346 | 30 | 85 | No | 24 | Female |
| 27 | BNT162b2/BNT162b2/BNT162b2/Moderna Bivalent | 209 | 298 | 30 | 96 | No | 26 | Female |
| 28 | mRNA-1273/mRNA-1273/mRNA-1273/Moderna Bivalent | 213 | 280 | 30 | 92 | No | 38 | Female |
| 29 | mRNA-1273/mRNA-1273/mRNA-1273/Moderna Bivalent | 269 | 276 | 23 | 94 | No | 40 | Male |
| *BA.5 breakthrough* | | | | | | | | |
| 30 | mRNA-1273//mRNA-1273/mRNA-1273/BA.5 | 182 | 127# | 32* | 88* | Yes | 60 | Female |
| 31 | mRNA-1273//mRNA-1273/mRNA-1273/BA.5 | 306 | 264# | 25* | 94* | Yes | 50 | Female |
| 32 | BNT162b2/BNT162b2/BNT162b2/BA.5 | 241 | 301# | 30* | 91* | Yes | 41 | Female |
| 33 | BNT162b2/BNT162b2/BNT162b2/BA.5 | 265 | 237# | 27* | 82* | Yes | 50 | Female |
| 34 | BNT162b2/BNT162b2/BNT162b2/BA.5 | 279 | 330# | 18* | 85* | Yes | 38 | Female |
| 35 | BNT162b2/BNT162b2/BNT162b2/BA.5 | 249 | 286# | 25* | 96* | Yes | 25 | Male |
| 36 | BNT162b2/BNT162b2/BNT162b2/BA.5 | 265 | 302# | 29* | 92* | Yes | 36 | Female |
| 37 | BNT162b2/BNT162b2/BNT162b2/BA.5 | 245 | 346# | 28* | 93* | Yes | 43 | Female |
| 38 | BNT162b2/BNT162b2/BNT162b2/BA.5 | 255 | 298# | 21* | 93* | Yes | 26 | Female |
| 39 | BNT162b2/BNT162b2/BNT162b2/BA.5 | 320 | 281# | 34* | 90* | Yes | 56 | Male |
| 40 | BNT162b2/BNT162b2/BNT162b2/BA.5 | 327 | 271# | 31* | 94* | Yes | 48 | Female |
| 41 | BNT162b2/BNT162b2/mRNA-1273/BA.5 | 312 | 285# | 18* | 84* | Yes | 55 | Female |

V2-V3 indicates days between vaccine doses two and three.
V3-V4 indicates days between vaccine doses three and four.
#V3-I indicates days between vaccine dose three and breakthrough infection of BA.5.

### Table S2. Summary of clinical cohorts

| Characteristic | Monovalent booster (N=15) | Bivalent booster (N=14) | BA.5 Breakthrough (N=12) |
|---|---|---|---|
| **Sex -- no. (%)** | | | |
| Female | 14 (93.3%) | 11 (78.6%) | 10 (83.3%) |
| Male | 1 (6.7%) | 3 (21.4%) | 2 (16.7%) |
| Mean Age (range)--yr | 55.3 (49, 63) | 37.8 (24, 49) | 44.0 (25, 60) |
| Mean days post vaccination or infection--1st blood draw (range) | 23.3 (20, 29) | 26.4 (23, 30) | 26.5 (18, 34) |
| Mean days post vaccination or infection--2nd blood draw (range) | 92.7 (74, 115) | 94.2 (82, 111) | 90.2 (82, 96) |
| Mean days between vaccines 2 and 3 (range) | 272.3 (201, 337) | 235.5 (205, 275) | 270.5 (182, 327) |
| Mean days between vaccines 3 and 4 or vaccine 3 and BA.5 infection* (range) | 188.0 (122, 294) | 302.6 (276, 346) | 277.3 (127, 346)* |
| **First and second vaccine doses** | | | |
| Pfizer (BNT162b2) | 15 (100.0%) | 10 (71.4%) | 10 (83.3%) |
| Moderna (mRNA-1273) | 0 (0.0%) | 4 (28.6%) | 2 (16.7%) |
| **Third vaccine dose** | | | |
| Pfizer (BNT162b2) | 15 (100.0%) | 8 (57.1%) | 9 (75.0%) |
| Moderna (mRNA-1273) | 0 (0.0%) | 6 (42.9%) | 3 (25.0%) |
| **Fourth vaccine dose** | | | |
| Pfizer | 15 (100.0%) | 3 (21.4%) | - |
| Moderna | 0 (0.0%) | 11 (78.6%) | - |

*For the BA.5 breakthrough cohort, days are measured between vaccine 3 and documented breakthrough infection.

### Supplementary References

1. Simon V, Kota V, Bloomquist RF, et al. PARIS and SPARTA: Finding the Achilles' Heel of SARS-CoV-2. mSphere 2022;7:e0017922.
2. Wang Q, Iketani S, Li Z, et al. Alarming antibody evasion properties of rising SARS-CoV-2 BQ and XBB subvariants. Cell 2023;186:279-86 e8.
3. Liu L, Iketani S, Guo Y, et al. Striking antibody evasion manifested by the Omicron variant of SARS-CoV-2. Nature 2022;602:676-81.
4. Iketani S, Liu L, Guo Y, et al. Antibody evasion properties of SARS-CoV-2 Omicron sublineages. Nature 2022;604:553-6.
5. Wang Q, Guo Y, Iketani S, et al. Antibody evasion by SARS-CoV-2 Omicron subvariants BA.2.12.1, BA.4, & BA.5. Nature 2022.
6. Liu L, Wang P, Nair MS, et al. Potent neutralizing antibodies against multiple epitopes on SARS-CoV-2 spike. Nature 2020;584:450-6.
